# ClgR contributes to pulmonary pathology but not bacterial growth in *Mycobacterium tuberculosis* infection

**DOI:** 10.1101/2020.02.04.934901

**Authors:** Uma Shankar Gautam, Deepak Kaushal

## Abstract

**Background:** The Clp proteases regulator, ClgR, is encoded in the *Mycobacterium tuberculosis* (Mtb) genome by Rv2745c gene *(clgR).* ClgR is required to clear damaged proteins, thereby preventing their accumulation in the cell. It also controls the availability of key enzymes or regulators via conditional degradation mechanism of proteolytic activity in Mtb [1,2].

**Methods:** It has been previously reported that Mtb *clgR* gene is induced in a sigma factor SigH-dependent manner and a deletion mutant of *clgR* is susceptible to growth in a hypoxic environment. Whether hypoxia is indeed a restriction factor and ClgR is required for Mtb growth in that environment remains unelucidated. We began to address this hypothesis in the C57/BL6 mouse model of TB where Mtb infected lungs do not form granuloma and the lung environment is considerably non-hypoxic.

**Results:** Our results demonstrate that despite not having a deficit in growth in either murine lungs or primary macrophages, in comparison to wild type, the *ΔclgR* mutant failed to induce pulmonary pathology.

**Conclusion:** We propose that ClgR is required for the pathogenesis of Mtb.

## Introduction

ClgR (Clp gene regulator) has been shown to be induced in response to stresses such as hypoxia, redox stress, and cell wall damages [3–8]. Proteolysis mechanisms regulated by ClgR are required for the resuscitation following the hypoxic shutdown of transcription activation mechanism and serve as a key phenomenon in the posttranscriptional regulation of gene expression in Mtb [9, 10]. Our previous findings suggest that the isogenic *ΔclgR* mutant is susceptible for growth during hypoxic stresses in-vitro [8]. The Mtb encoded Clp protease system is essential for the degradation of potentially toxic unwanted proteins and prevents their accumulation in the cell [10–14]. Mtb encounter multitude of host pressure within macrophage and other lung cell types during host-pathogen interactions. As a result, Mtb experiences extensive damage to cellular proteins and their removal is crucial for the survival of the pathogen. The robust induction of *clgR* during hypoxia [7] clearly indicates that Clp gene regulator could also be critical within intragranolomatous environment where most bacilli persist and encounter host pressures such as hypoxia and redox stress [7, 8, 15–17]. However, we do not know what targets are proteolyzed during infection, and whether the spectrum of Clp targets is altered by stresses in-vivo. Here, we show that *ΔclgR* is attenuated but not compromised in growth in mice. We propose to study the ClgR as control system in host-pathogen interactions via the post-transcriptional mechanisms of targeted proteolysis in future studies.

## Materials and Methods

### Ethics statement

Tulane National Primate Research Center (TNPRC) facilities are accredited by the American Association for Accreditation of Laboratory Animal Care (AAALAC) and licensed by the U.S. Department of Agriculture. The animals are usually group-housed in appropriate social settings in accordance with the guidelines of AAALAC, which annually inspects all facilities. The guidelines prescribed by the National Institutes of Health Guide to Laboratory Animal Care were routinely followed for animal care. The Humane endpoints were applied as a measure of reduction of discomfort. All the procedures used in this study were from an approved protocol of Institutional Animal Care and Use Committee and the Institutional Biosafety Committee (IACUC). The IACUC performs semiannual and annual inspections of the facility to certify compliance with the highest possible levels of housing conditions, environmental enrichment and feeding regimens. Experiments involving *Mycobacterium tuberculosis* Mtb and *ΔclgR* strain (used in this study were approved by the Tulane Institutional Biosafety Committee.

### Mice and Aerosol Infection

*Mycobacterium tuberculosis* wild type strain CDC1551 (henceforth referred to as Mtb) and an isogenic deletion mutant of *Rv2745c, Mtb:ΔclgR* [7] henceforth referred to as *ΔclgR* were cultured as previously described [15, 18]. Two groups of 56 male C57/BL6 mice (5-6 wk old) (Jackson Laboratory, Bar Harbor, ME) were infected with approximately 50 CFUs via aerosol [15] (Table 1). Following the day of infection, four mice were euthanized to determine the bacterial deposition in lung and thereafter four mice per group were killed every 4 weeks for CFU count and lung histopathology (day1, week 4, 8, 12, 16, 20, and 50) as described [15].

**Table 1.**
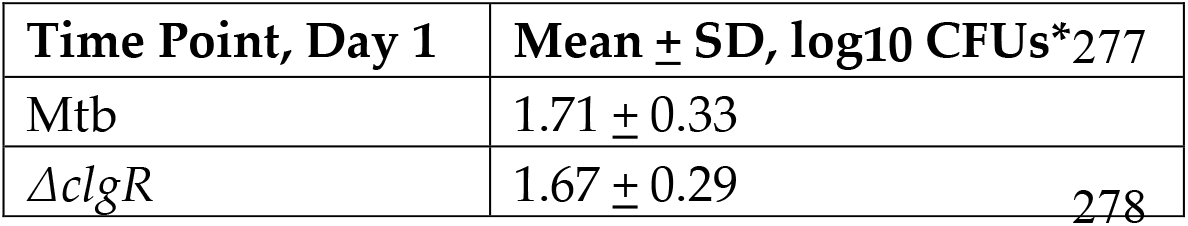
Pulmonary Bacterial Burdens in C57/BL6 Mice after Aerosol Infection

### Staining Procedures, Histopathology, and Microscopy

Lung tissue was fixed for hematoxylin-eosin histology and confocal microscopy strictly following the detailed experimental procedures described previously [15, 16, 19–23].

### Isolation of Bone-Marrow derived macrophages and Infection with Mtb, *ΔclgR,* and a complemented strain

Isolation and maintenance of Rhesus Bone Marrow Derived Macrophages (Rh-BMDMs) as well as their infection with Mtb, isogenic mutant *ΔclgR* and a complemented strain was carried out strictly as per the experimental details described previously [21]. Briefly, Rh-BMDMs were cultured in IMDM complete media (Gibco) (IMDM media supplemented with 10% heat-inactivated FBS (Hyclone) and 1% Penicillin/Streptomycin mix (Pen/Strep, Gibco). Rh-BMDMs were infected at a multiplicity of infection (MOI) of 10:1 (10 bacteria per 1 cell). The infected cells were lysed (0.1% saponin) for CFU assay up to 72 hr post-infection.

### Statistics

Statistical analyses used GraphPad Prism 6.0b (GraphPad Software, La Jolla, CA).

## Results

### Deletion of *clgR* does not alter the growth phenotype of *Mycobacterium tuberculosis* in C57/BL6 mice

We tested the infection phenotype of *ΔclgR* in comparison to wild type Mtb by comparing the growth in mice lung and spleen. The CFU data indicates that *ΔclgR* is not compromised for the infection (Table 1) or growth in mice (Fig. 1). In comparison to Mtb, significantly higher levels of *ΔclgR* CFU were detected in lung tissue (Fig. 1). To rule out any defect in the occurrence of bacterial burdens during the dissemination process, we measured the bacterial burdens present in spleen in all the infected mice at various time points throughout the study (Fig. 1). While at week 4, we observed a lower bacterial in mice infected with mutant *ΔclgR,* the mutant was able to catch up in and generally maintained equivalent or higher burden in this tissue than the Mtb infected mice. Thus, the CFUs in the mice infected with *ΔclgR* were ~2.5 fold, ~3 fold, ~2 fold higher in mice lung at week 8, 12 and 16 in comparison to Mtb. At week 20 and 50, CFUs of *ΔclgR* in spleen were indistinguishable from Mtb.

**Fig. 1.**
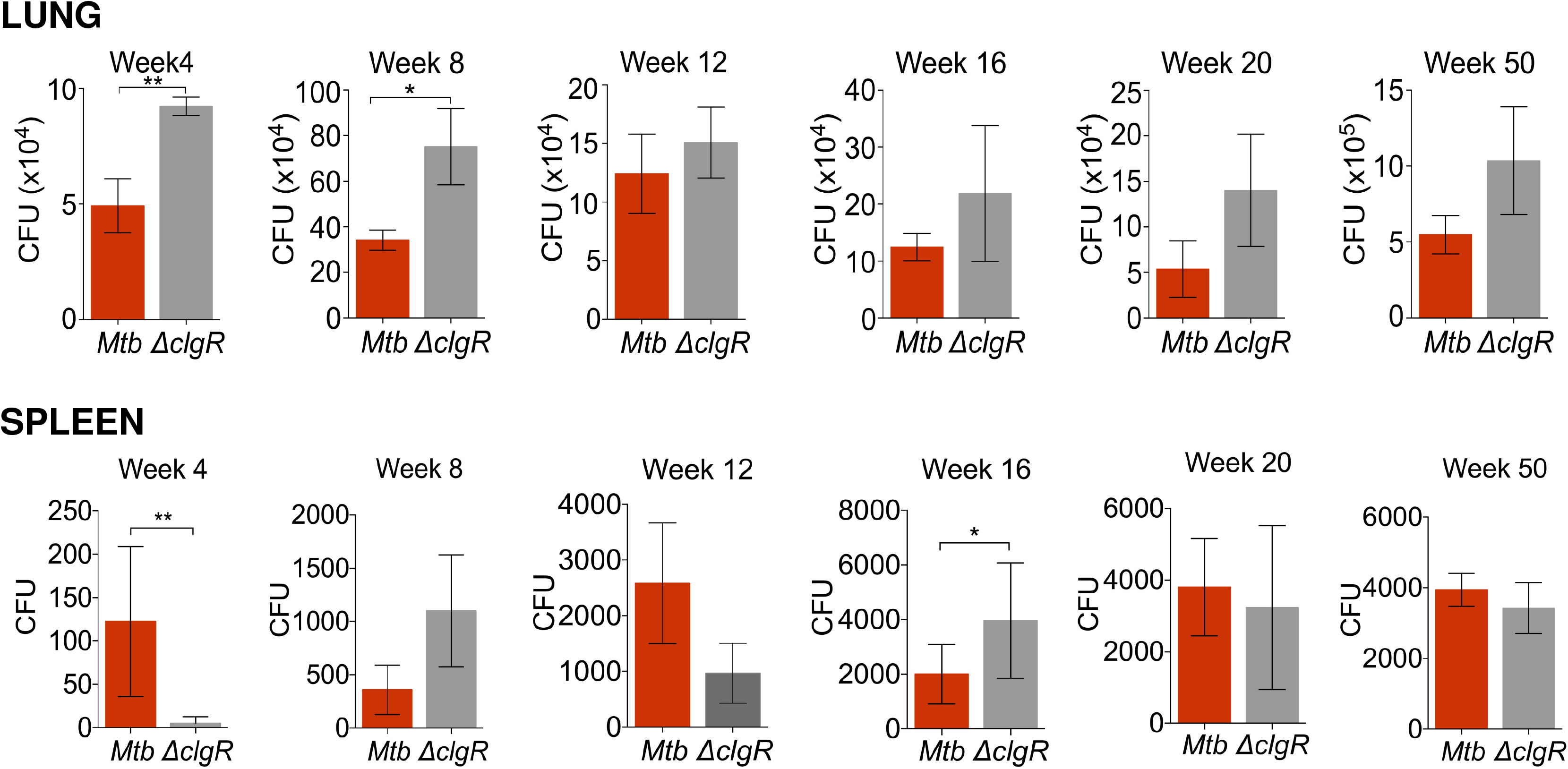
Measurement of bacterial growth in mice. The bacterial burdens determined in mice lung (top panel) and spleen (lower panel) at weeks 4-50; Mtb (red), *ΔclgR* (grey). Results are expressed as CFUs in the entire tissue based on weight. *P < 0.05; **P < 0.005 (unpaired *t* test).

The CFU data in spleen demonstrates that the growth phenotype of the *ΔclgR* is not altered during the process of dissemination. Hence, the survival phenotype of *ΔclgR* in the spleen was very similar to that of lung. The CFU data demonstrate that bacterial burdens in mice infected with the *ΔclgR* were maintained at a level at least equal or were higher ~2-3 fold than Mtb at each time point (Fig. 1). We noted the comparable bacterial numbers between two groups except at week 4 and 16 in spleen where bacterial numbers were statistically significant at these timepoints between two groups (P <0.05) (Fig. 1). These results clearly indicate that *ΔclgR* is neither compromised for growth (Fig. 1) nor establishment of infection in C57BL/6 mice (Table 1). Next, we measured the involvement of mice lung in disease progression by comparing the lung histopathology during the course of infection between two groups.

### Lung histopathology measurement in C57/BL6 mice

We measured the disease progression in infected mice at week 4, 8, 12, 16, 20, and 50 by histopathology. Despite comparable or higher lung CFUs, mice infected with *ΔclgR* exhibited lower lung involvement in comparison to Mtb group throughout the course of infection, and the differences were statistically significant at later stages (weeks 16, 20 and 50) (Fig. 2). For example, at week 16, lung involvement of the *ΔclgR* bacteria was 3% compared to Mtb (25% lung involvement) and this involvement was ~10% in *ΔclgR* bacteria in comparison to Mtb (~33% lung involvement) by week 50 (Fig. 2). Together, these results demonstrate that despite comparable to higher bacterial burdens in the lung, *ΔclgR* fails to induce comparable pulmonary pathology in comparison to Mtb. We therefore conclude that *ΔclgR* is not compromised for growth in mice but causes reduced immunopathology (Fig. 2). Clearly the minimal pathology observed in the *ΔclgR* group is inconsistent with the cognate bacterial. Next, we determined the growth phenotype of *ΔclgR* and Mtb in primary macrophages in vitro.

**Fig. 2.**
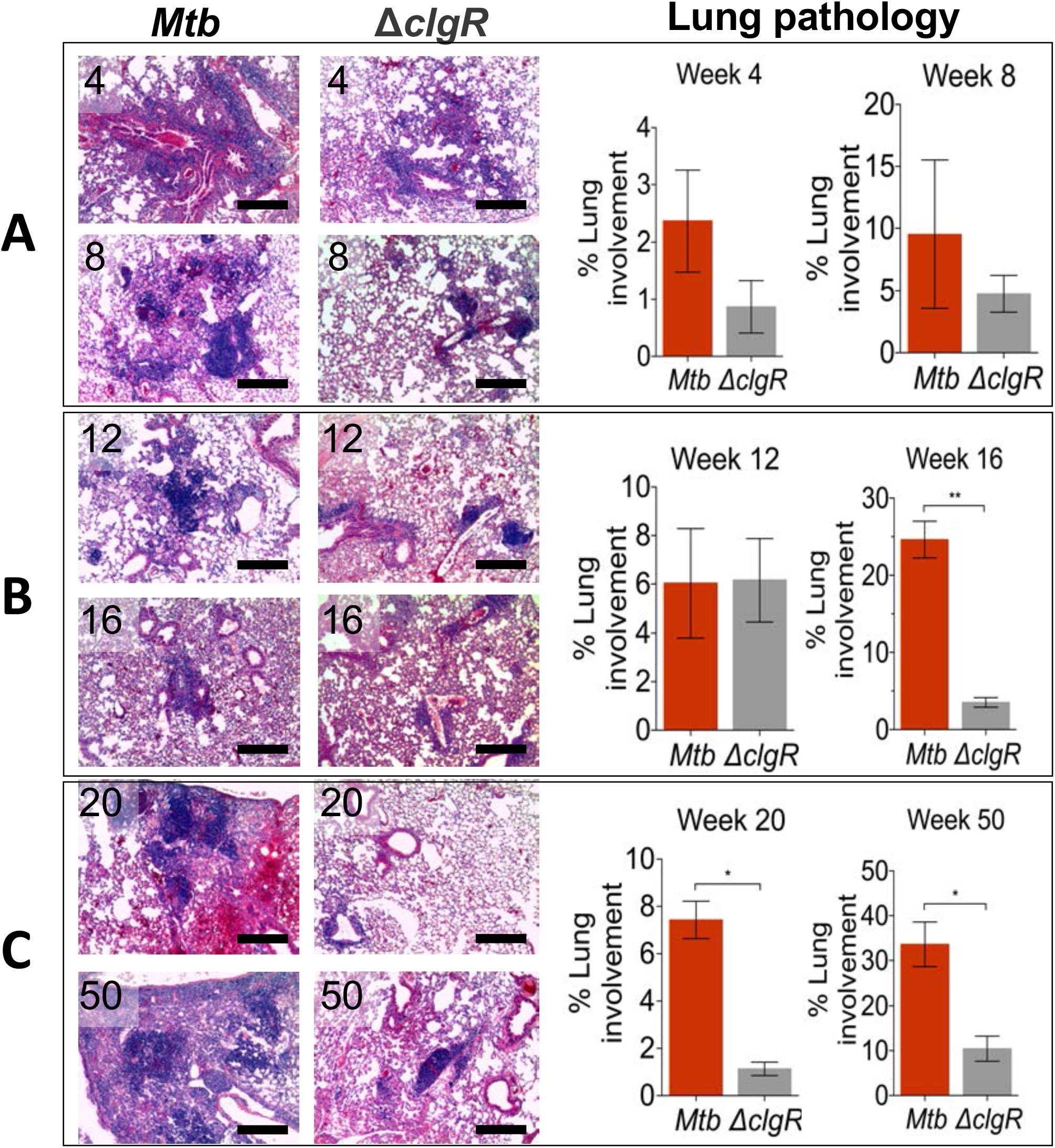
Lung histopathology in infected mice. Haematoxylin and eosin (H&E) staining from representative animals (lefts panels A-C) and their lung pathology scores shown as percentage of lung area grouped from lung lobes of four mice in both Mtb (red) and *ΔclgR* (grey) groups at weeks 4-50 (right panels A-C). Lung areas with multiple inflammatory lesions were present in Mtb groups versus fewer in ΔclgR-infected animals (panels A-C). Significant differences in lesion scores between two groups at the latter time point. The data are means±sd, *P < 0.05; ** *P* < 0.005 (unpaired *t* test). Black scale bars in (a–c), 25 μm.

### Growth Phenotypes in primary macrophages

Next, we evaluated the growth pattern of mutant *ΔclgR* in primary macrophages derived from rhesus macaques, a highly human-like model of TB pathogenesis. The growth of *ΔclgR* was compared up to 3 days in comparison to wild type Mtb and a complemented strain of *clgR* as a correlate of protection from TB. CFU results demonstrated that the mutant *ΔclgR* did not exhibit growth restriction in RhBMDMs as it replicates comparable to wild type Mtb as well the complement strain in a time-dependent manner (Fig. 3).

**Fig. 3.**
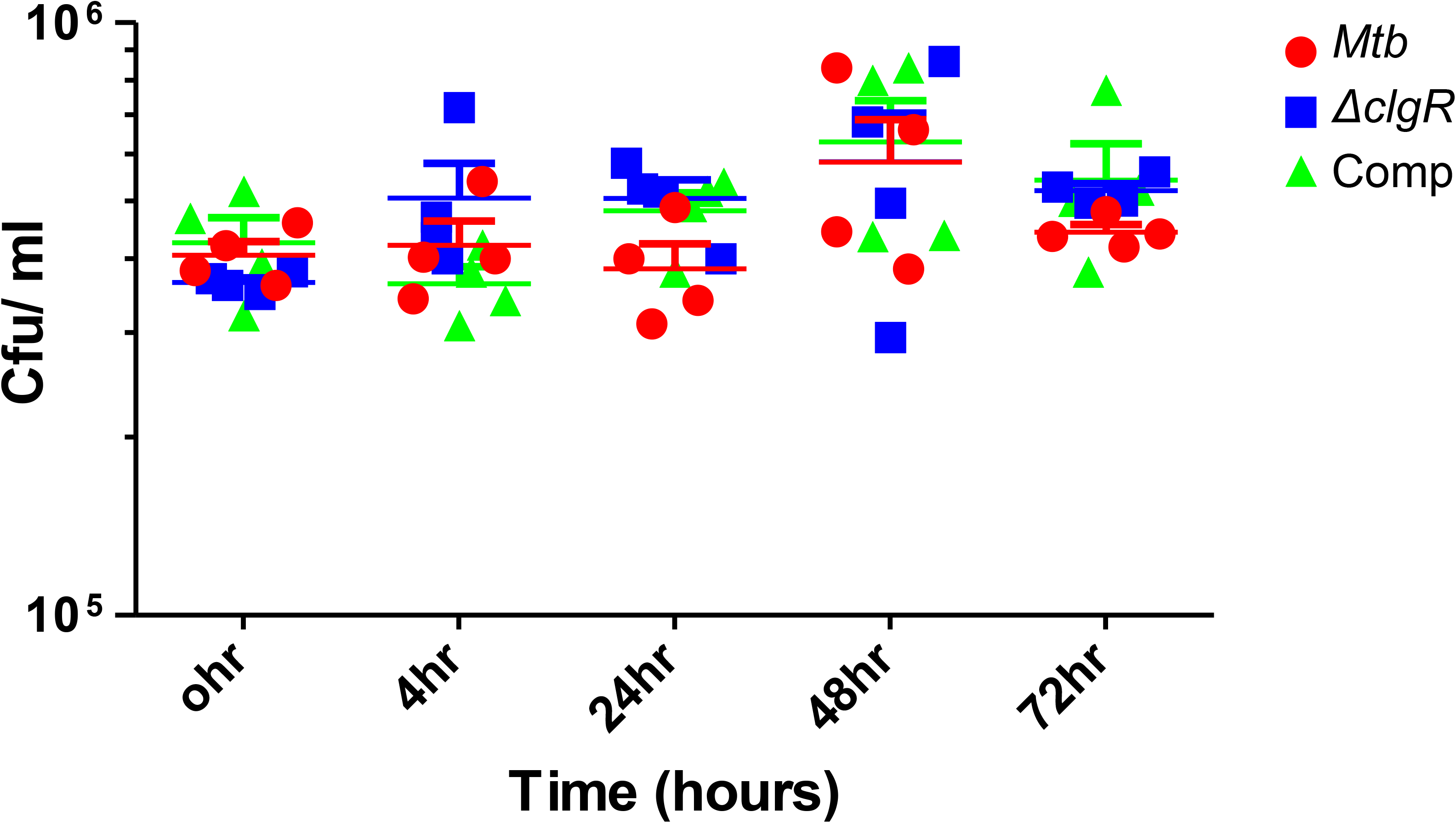
Comparative measure of bacterial growth. Rhesus macaque BMDM (RhBMDM) *in-vitro* killing assay in CFU ml^-1^ with Mtb (red), *ΔclgR* (blue), and Complemented strain (green). The data (means±sd) are Statistically insignificant using two-way ANOVA with multiple comparisons.

## Discussion

We have previously reported that the *ΔclgR* mutant is compromised in growth when cultured in inducing stress conditions such as hypoxia or redox environment in vitro [7, 8]. In murine lungs and spleens, bacterial burdens of *ΔclgR* were not any lower than Mtb, in fact, at some time points significantly more *ΔclgR* bacilli could be isolated from murine tissues. The higher bacterial replication rate of the *ΔclgR* mutant in C57BL/6 mice (Fig. 1) could mean that ClgR and the products of its regulation are critical for in vivo survival of Mtb [7, 8]. However, since our studies were conducted in C57/BL6 mice, where lesions are not known to be hypoxic, our results do not insinuate ClgR as being critical for survival during hypoxia in vivo. Hypoxic models such as Kramnik mice [24, 25] or macaques [17, 22] would be required to address those questions. Our mouse data collectively supports the idea that the *clgR* deletion neither limits the ability of Mtb to grow in the lungs infection nor reactivation to extrapulmonary tissues. We further verified this growth phenotype in infected primary macrophages and determined that similar growth phenotype pattern was observed in vitro (Fig. 3.) McGillivray *et al.* have previously demonstrated this mutant was defective for growth in standing flasks following hypoxic treatment [8]. This may be due to non-stimulation of DosRST regulon, and/ or due to an accumulation of hypoxia in a time-dependent manner [7, 15, 17].

The key result from our studies is that despite being able to grow at least equally well if not better than Mtb, the *ΔclgR* mutant resulted in significantly reduced pulmonary pathology (Fig. 2). This result is not surprising. A mutant in the master regulator of redox stress SigH *(AsigH)* also exhibits a comparable phenotype [26]. ClgR is responsive to redox stress and induced by it in a sigH-dependent manner ([7, 8]. Therefore it is conceivable that the reduced pathology despite bacterial burden phenotype exhibited by the *ΔclgR* mutant is derived from oxidative stress in murine lungs. While it remains to be determine whether the infection phenotype of *ΔclgR* can also be elicited and provide protection against TB in human-like macaque model [15, 17, 22, 27, 28] and C3HeB/FeJ mice [15, 25] where granulomas are hypoxic and may virtually not only limit the growth of *ΔclgR* but may also result in complete clearance of TB infection [15–17, 19, 20, 22]. While it remains to be elucidated, our data suggest that *ΔclgR* may elicit highly protective responses against TB in macaques.

